# FORGE audits residue-level information encoded in RNA tertiary-structure geometry

**DOI:** 10.64898/2026.07.05.736550

**Authors:** Leighton Gow, Jintang Li, Xianhan Tan, Ke Liang, Ning Gui, Binli Luo

## Abstract

Coarse RNA coordinate representations are widely used, yet the biological information they encode remains unquantified. We introduce FORGE, which converts a seven-atom RNA geometry representation into 935 interpretable descriptors and reports which residue-level annotations this geometry supports. On 4,135 post-2025 RNA chains, FORGE recovered 64.6% of native nucleotides; a six-atom control lacking the glycosidic nitrogen retained 58.5%, locating most of this signal in phosphate–sugar geometry. Confidence was sharply graded: abstaining from the least-confident half of positions raised accuracy to 94.4%, yet many chains remained only partially identifiable. The same descriptors predicted base-pair state far better than a DMS-like proxy or protein-proximal context. Native–decoy, OpenKnot and solved-pseudoknot analyses showed that nucleotide identifiability, foldability and experimental design score are separable: AlphaFold 3 reproduced the experimental fold for one of four AI-designed constructs and none of the sequences FORGE read from their geometry. FORGE provides a reproducible audit layer for RNA structural interpretation.

---

RNA structures are increasingly resolved by crystallography, cryo-electron microscopy and NMR, with recent workflows extending cryo-EM structure determination to RNA-only systems [1, 2]. In parallel, geometric learning and inverse-folding models now generate, score and compare RNA folds at scale [3–6]. Yet a basic question remains under-specified: what information is actually present in a coarse RNA tertiary-structure representation? A phosphate–sugar backbone with a glycosidic anchor encodes local geometry, stacking, helical organisation and packing, but omits base-edge chemistry, modifications, solvent, ligands, proteins and conformational ensembles. Reading such a representation as if it fully specified sequence, foldability or function therefore risks overinterpreting the coordinates. Structural biologists and modellers need to know whether a residue assignment or a designed sequence is supported by the deposited backbone, or whether additional evidence, such as density, covariation, chemical probing or full-atom refinement, is required.

This issue is clearest in RNA inverse folding, where models predict nucleotide sequences from target three-dimensional backbones and are commonly evaluated by sequence recovery, the fraction of positions at which a predicted nucleotide matches the deposited native sequence [4–6]. Recovery is useful but conflates several properties. A deposited nucleotide may be identifiable because the local backbone geometry sharply constrains base identity. Alternatively, the same geometry may tolerate several nucleotides, with the native base reflecting evolutionary history, ligand context or functional selection rather than a unique structural requirement. Sequence recovery therefore mixes structural identifiability, native-like compatibility and model behaviour, and is not by itself evidence of generative design success.

A more useful view treats RNA tertiary structure as an information source with multiple measurable layers. Some annotations, such as nucleotide identity or base-pairing state defined from three-dimensional geometry [7], lie close to the backbone. Others, such as chemical-probing reactivity or protein-binding context, depend on nucleotide chemistry, solvent exposure and the external molecular environment [8]. A computational tool that maps these layers, reports uncertainty and exposes where the representation is underdetermined would complement both sequence design and manual structure interpretation.

Here we introduce FORGE (Feature-engineered RNA Geometry Evaluation), which distils RNA tertiary structure into 935 explicit, interpretable geometric descriptors from a seven-atom backbone representation. Gradient-boosted trees trained on these descriptors report per-position nucleotide probabilities, confidence scores and sequence–backbone compatibility scores [9], and the same representation supports base-pair prediction, a RibonanzaNet-derived DMS-like proxy and protein-proximal context. The backbone representation thereby becomes an auditable set of task-specific readouts rather than a single recovery score. The goal is not universal RNA design capability, but a quantification of which annotations the geometry supports, when the model should abstain and where additional evidence is required.

We evaluate FORGE under a temporal split of Protein Data Bank RNA structures [1], training on pre-2025 depositions and testing on 4,135 post-2025 chains.

The evidence ladder follows the intended use: temporal sequence identifiability, representation controls, confidence and abstention, task-specific readouts, native-versus-decoy compatibility and stress tests on solved AI-designed pseudoknots. Together, these analyses show that the seven-atom backbone carries substantial but bounded nucleotide-identifying information, most of which is retained without the glycosidic nitrogen coordinate, and that structural identifiability, foldability and experimental design score are distinct quantities.

## 1 Results

### 1.1 FORGE reports residue-level probabilities from explicit backbone features

FORGE takes an RNA structure in PDB or mmCIF format, extracts a seven-atom representation per nucleotide (P, O5′, C5′, C4′, C3′, O3′ and N1/N9), builds a *k* = 32 nearest-neighbour graph and computes 935 geometric descriptors (Fig. 1): 211 node features plus 724 edge-derived features pooled to residues. A multi-class XGBoost classifier then reports probabilities for A, C, G and U, together with a confidence score. The same output yields a cross-entropy compatibility score for any candidate sequence and position-level SHAP explanations on request [10]. The intended output is not only a sequence call but a diagnostic statement about which positions are identifiable from geometry and which require orthogonal evidence.

**Fig. 1.**
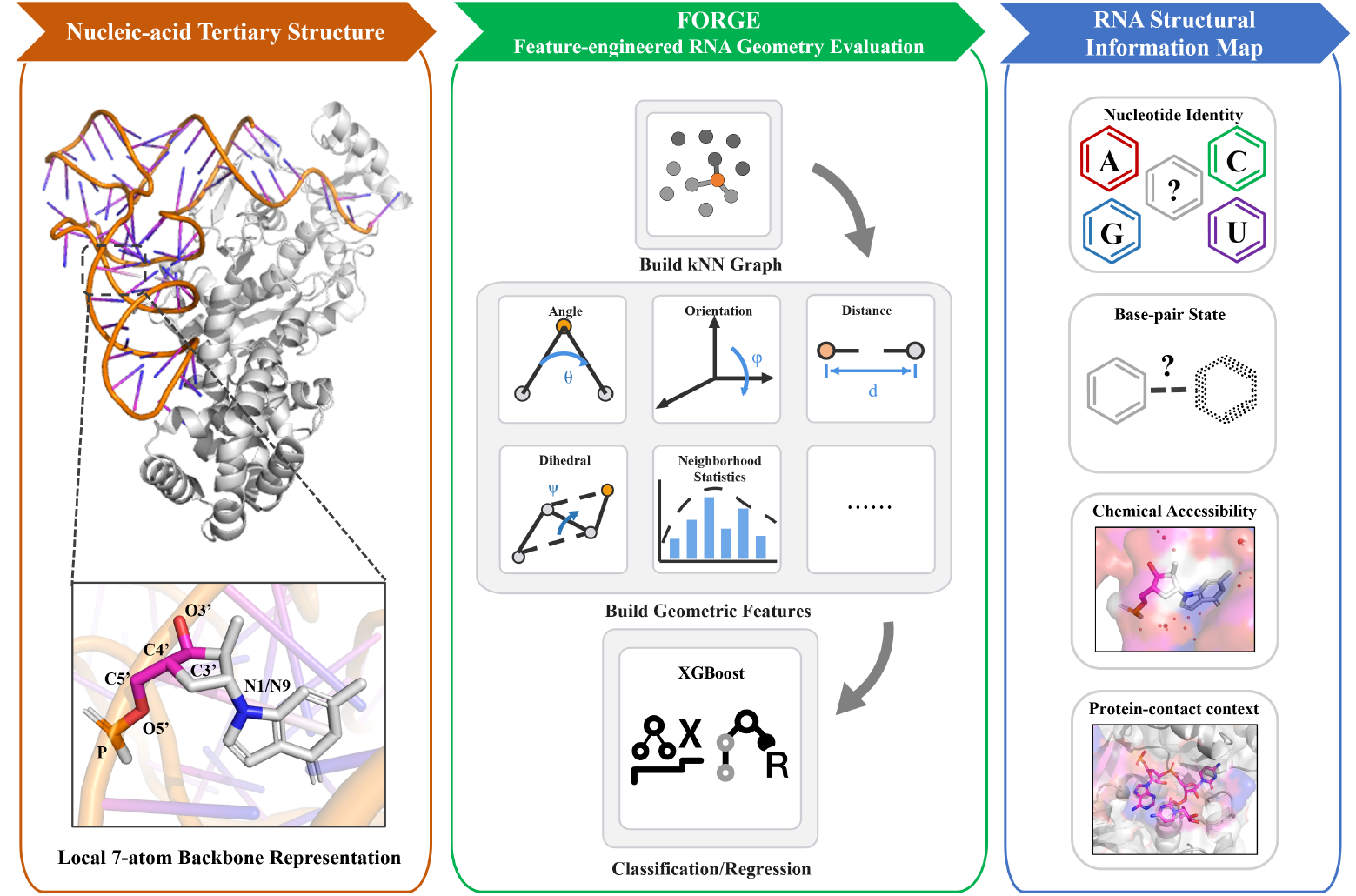
FORGE provides a calibrated audit workflow for RNA tertiary structures. RNA structures are reduced to a seven-atom geometry representation, converted into a residue-neighbour graph and featurised into 935 interpretable descriptors. Task-specific models then report nucleotide probabilities, confidence, sequence–backbone compatibility and auxiliary readouts for base-pair state, a DMS-like proxy and protein-proximal context, identifying which residues are geometrically constrained and which require additional chemical, evolutionary or experimental evidence.

### 1.2 Temporal testing shows bounded nucleotide identifiability

We trained FORGE on 13,251 pre-2025 RNA chains and evaluated it on all 4,135 post-2025 chains in the locked manifest (291,331 nucleotides). Weighted recovery was 64.6% (95% CI [63.7, 65.6]%, chain-level bootstrap) and mean per-chain recovery was 63.4% (Fig. 2a,b). An MLP trained on the same 935 features reached 59.5%, showing that most of the signal resides in the explicit geometric representation rather than in the tree ensemble.

**Fig. 2.**
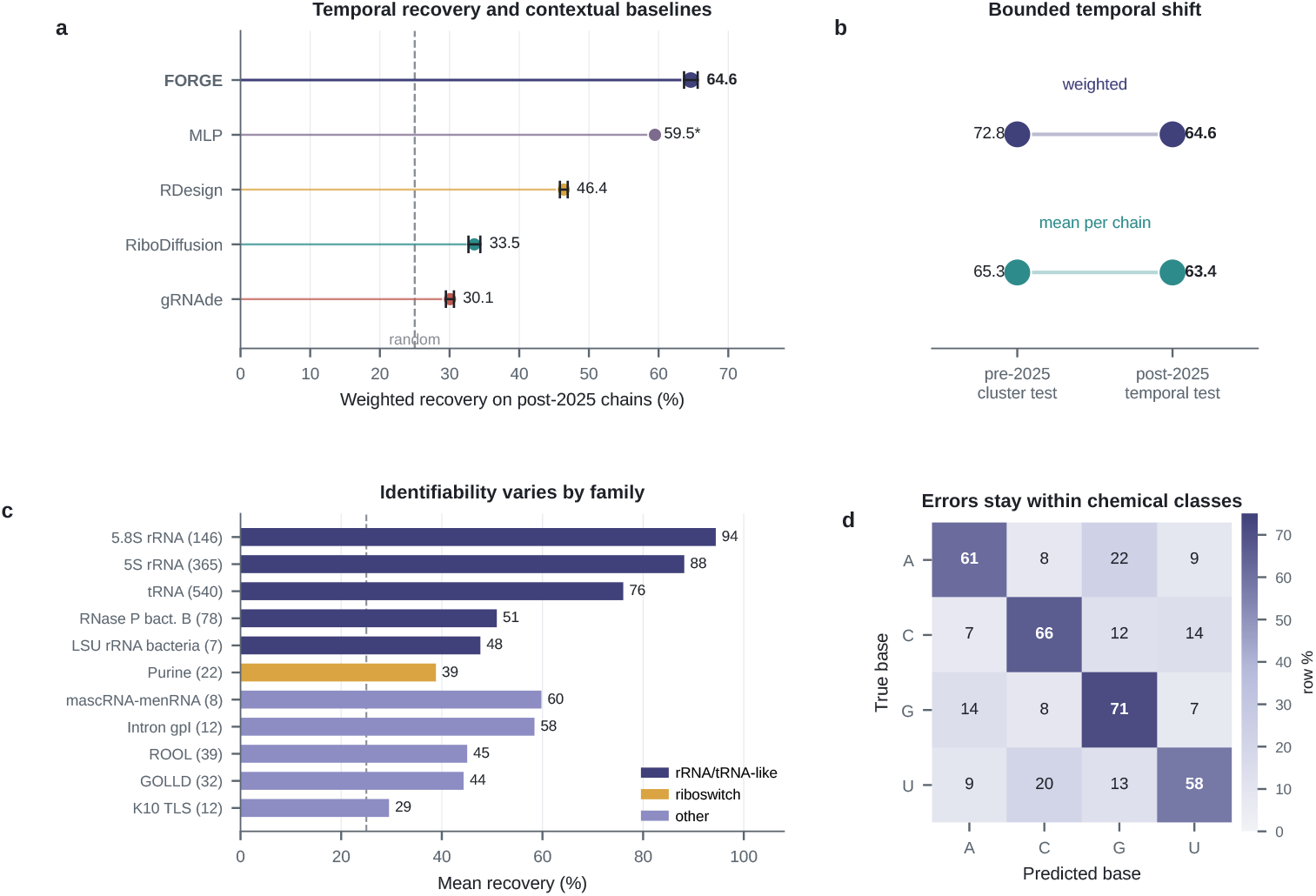
Nucleotide identity is partially readable from backbone geometry, but unevenly across families and bases. **a**, Weighted recovery of FORGE, an MLP trained on the same features and public pretrained baselines on post-2025 RNA chains (horizontal lollipops; black ticks, chain-bootstrap 95% intervals where per-chain prediction artifacts were available; the asterisk marks the MLP point estimate, for which no per-chain artifact was stored; dashed line, random recovery). This panel is not a same-training benchmark; RDesign was evaluable on fewer chains (3,966) than the other methods (4,135). **b**, FORGE recovery on the pre-2025 cluster-held-out test and the post-2025 temporal test, for length-weighted and mean per-chain recovery. **c**, Mean per-family recovery for Rfam-labelled post-2025 chains, coloured by broad RNA class, with chain counts in parentheses; several families have small sample sizes. **d**, Row-normalised confusion matrix for the temporal per-position predictions; values are percentages within each true nucleotide class.

For context, we evaluated public inverse-folding models on the same post-2025 structures using their pretrained outputs [4–6]. Weighted recovery was 30.1% for gRNAde (2.1M parameters), 33.5% for RiboDiffusion (52.2M) and 46.4% for RDesign (~1.5M) on the chains for which each baseline was evaluable. These models differ in training data, architecture and input atom set and were not retrained on the FORGE split, so they orient the reader rather than constitute a controlled benchmark.

Recovery varied strongly by structural family. Regular ribosomal and tRNA families were easiest (5.8S rRNA, 94.4%; 5S rRNA, 88.2%; tRNA, 76.0%), whereas several riboswitch and pseudoknot-rich families were much harder (Purine, 38.8%; K10_TLS, 29.5%; Fig. 2c). Error patterns were chemically interpretable: G had the highest diagonal recovery (71.1%), while U was least recovered (57.8%) and most often confused with C (Fig. 2d), consistent with a representation that carries strong local geometric signal but lacks the base-edge atoms needed to resolve A/G and U/C distinctions.

### 1.3 Most nucleotide-discriminating signal remains without the glycosidic nitrogen

To locate the origin of the nucleotide signal, we first aggregated the existing model’s four-class output into P(purine) and P(pyrimidine), which classified chemical class at 85.1% accuracy (random 50%). We then physically removed the N1/N9 coordinate before feature extraction and retrained from scratch on the full pre-2025 training set. On the post-2025 temporal test, this six-atom (N-free) model retained 58.5% weighted recovery (95% CI [57.4, 59.6]%), only 6.2 percentage points below the seven-atom model, and 72.4% purine/pyrimidine accuracy (Fig. 3a). A matched-subset retraining control gave consistent results (69.9% versus 66.7%; Supplementary Fig. S3). Most of the purine/pyrimidine signal therefore resides in phosphate–sugar geometry itself.

**Fig. 3.**
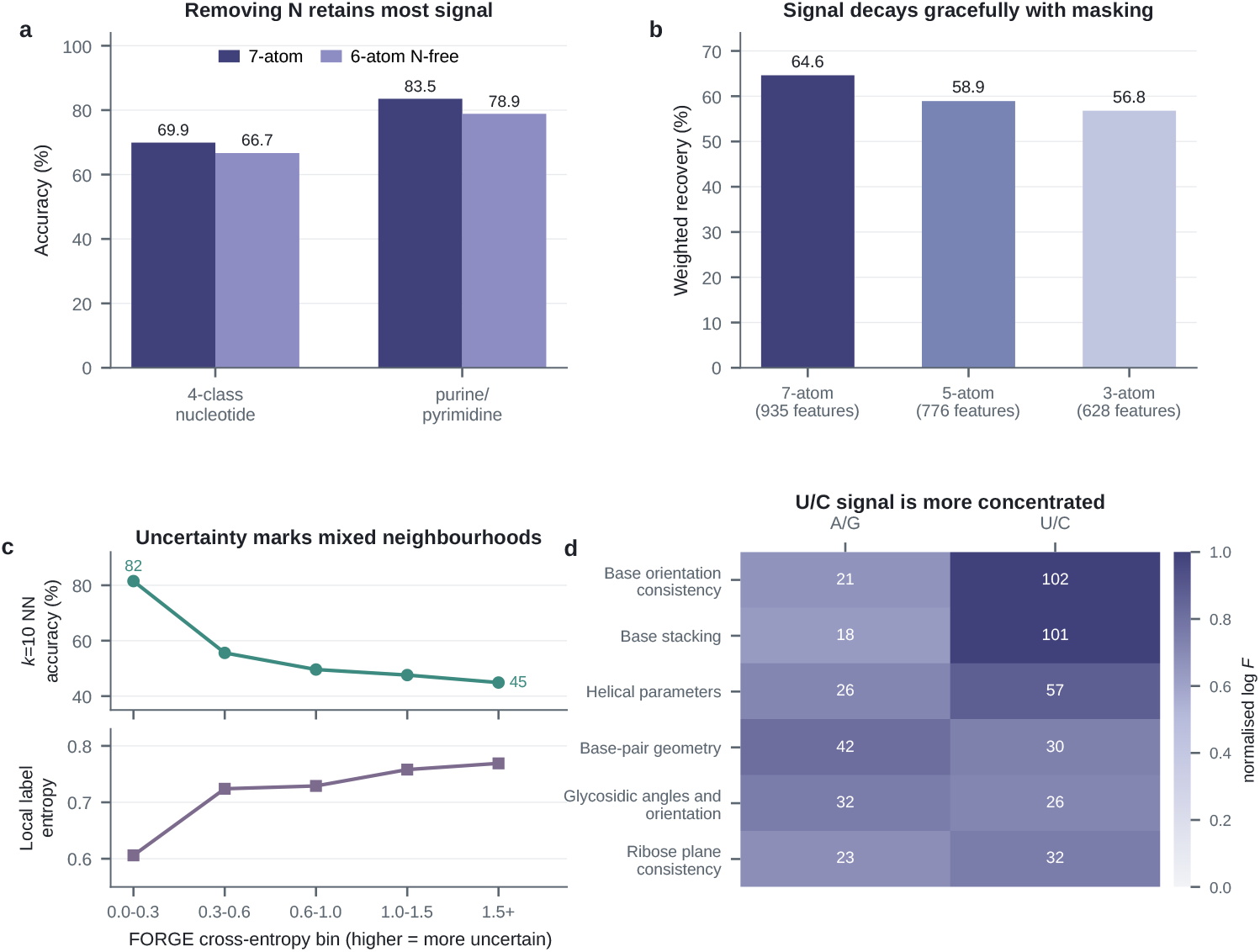
Representation controls and feature-space ambiguity. **a**, Matched-subset retraining after removing the glycosidic nitrogen coordinate before feature extraction (4-class recovery and purine/pyrimidine accuracy; the corresponding full post-2025 temporal comparison is 64.6% versus 58.5%). **b**, Inference-time atom-feature masking on the full post-2025 set (7-, 5- and 3-atom masks with the number of retained features). **c**, Leave-one-chain-out *k* = 10 nearest-neighbour accuracy (top) and local label entropy (bottom) across FORGE cross-entropy bins. **d**, Feature-ontology categories with the strongest discrimination for the A/G and U/C ambiguous pairs (colour, normalised log-transformed ANOVA *F*-statistic; numbers, raw *F*).

Inference-time atom-feature masking showed the same graceful decay: weighted recovery moved from 64.6% with all seven atoms to 58.9% after removing O5′/O3′-dependent features and to 56.8% when only P-, C4′- and N-dependent features remained (Fig. 3b). These masks are sensitivity tests without retraining, whereas the retrained N-free result (58.5%) is the stronger evidence that phosphate–sugar geometry carries substantial nucleotide-discriminating information.

Feature-space analyses supported the same conclusion under a leave-one-chain-out design. On 1,134 held-out query positions, a *k* = 10 nearest-neighbour classifier in the 935-dimensional feature space matched XGBoost almost exactly (53.6% versus 53.9%). Across FORGE cross-entropy bins, *k* = 10 accuracy fell from 81.5% to 44.9% as local label entropy rose from 0.606 to 0.769 (Spearman *ρ* = +0.227; Fig. 3c): uncertain positions occupy feature-space regions with genuinely mixed nucleotide labels. A feature ontology covering 889 of 935 dimensions further showed that U/C discrimination concentrates in base-orientation and stacking features, whereas A/G discrimination is weaker and more distributed (Fig. 3d).

### 1.4 Confidence identifies high-accuracy subsets and chain-level failure modes

A diagnostic model must report when it should not be trusted. On the 2,584 post-2025 chains (143,743 positions) for which the full calibration analysis was available, FORGE had an expected calibration error of 0.055 and a pooled accuracy of 73.1% (Fig. 4a). Removing the least-confident 50% of positions raised retained-position accuracy to 94.4%, and removing 75% raised it to 99.5% (Fig. 4b). Consistent with this, errors concentrate at low confidence (Fig. 4c). The calibration subset is easier than the full temporal set, so its 73.1% pooled accuracy should not be substituted for the 64.6% full-set weighted recovery (Supplementary Table S1).

**Fig. 4.**
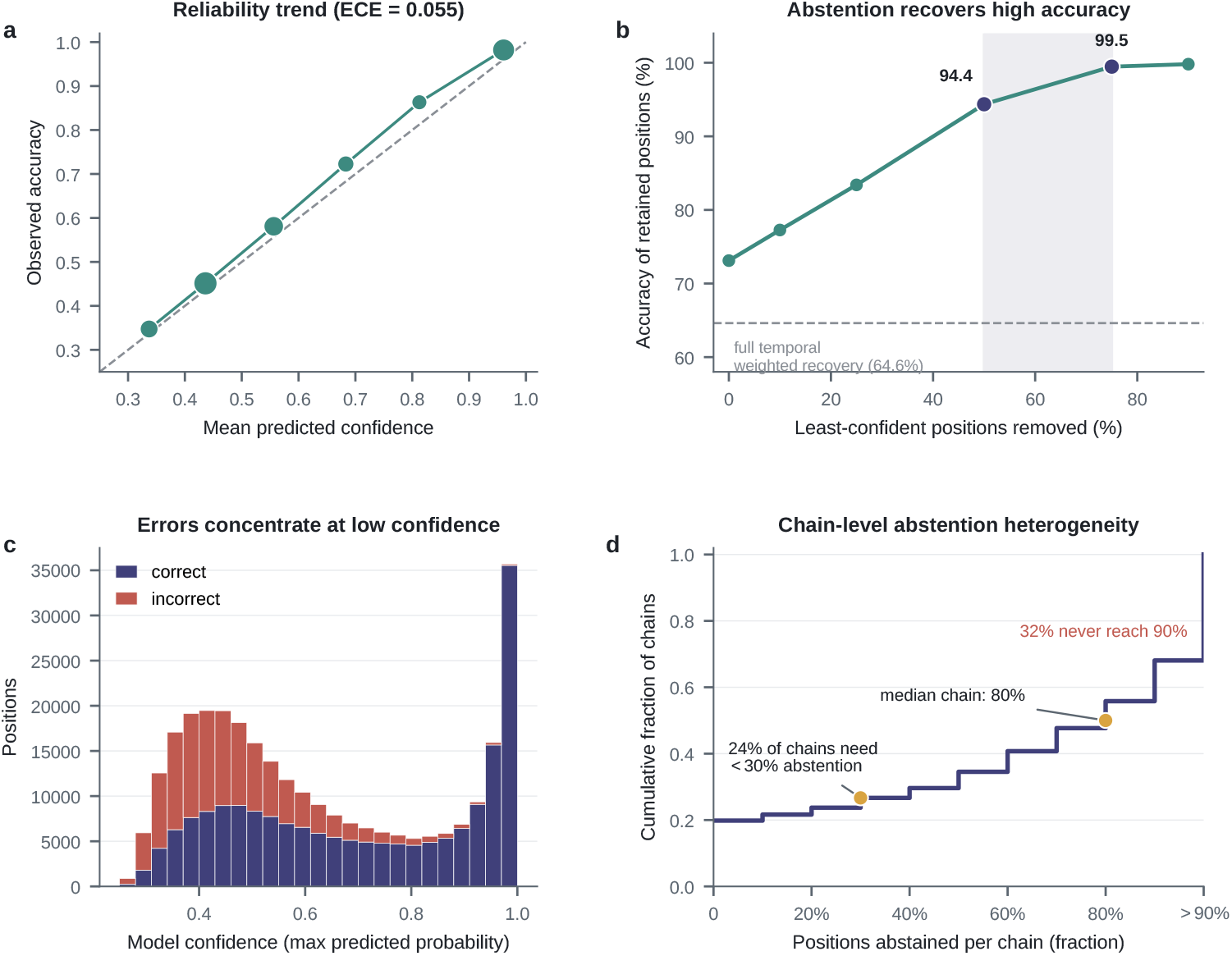
Confidence calibration and abstention. **a**, Reliability trend from retained post-2025 predictions (marker size, positions per bin); ECE was computed on the locked calibration-evaluable subset. **b**, Pooled abstention curve on the same subset; the dashed line marks full temporal weighted recovery and the shaded band marks the 50–75% abstention operating range. **c**, Stacked histogram of model confidence for correct and incorrect per-position predictions on the full temporal set. **d**, Empirical cumulative distribution of the per-chain abstention fraction required to reach 90% retained-position accuracy; chains in the *>*90% bin did not reach the target after removing 90% of positions.

Per-chain abstention gives a more conservative operational view. Across the full post-2025 set, the median chain required abstaining from 80% of its positions to reach 90% retained-position accuracy, 24% of chains needed fewer than 30% removed, and 32% did not reach the target even after removing 90% of positions (Fig. 4d). FORGE therefore supports high-confidence annotation where backbone geometry is sufficiently constraining, but for many chains the majority of positions are underdetermined by local geometry alone.

### 1.5 Backbone features support direct geometric labels more strongly than external context

If FORGE is more than a nucleotide classifier, the same features should contain different amounts of signal for different annotations. We therefore trained task-specific XGBoost models on the 935-dimensional representation for three further labels chosen to span direct geometry, partial backbone exposure and external environment: DSSR base-pair state [7], a RibonanzaNet-derived DMS-like proxy and protein-proximal context.

The expected ordering was observed (Table 1). Nucleotide identity showed the strongest four-class signal (64.6% weighted recovery; random 25%). Base-pair state, tightly coupled to A-form geometry, was predicted at 79.2% accuracy against a 63.1% majority baseline (AUC 0.880). The DMS-like proxy was captured at intermediate strength (Pearson *r* = 0.576, *R*^2^ = 0.329), consistent with reactivity depending on both local conformation and nucleotide chemistry in probing assays [8]. Protein-proximal context showed a weak but detectable signal (AUC 0.673, F1 0.461), as expected for a label that depends on protein surfaces and contacts not represented in the RNA-only backbone. This last task should be read as detecting geometry-associated signal correlated with protein proximity, not as residue-level binding-site prediction; performance may partly reflect structural differences between complex-forming and isolated RNAs.

**Table 1.** Task-specific readouts from the same backbone descriptors. All tasks use the same 935 geometric descriptors computed from RNA coordinates; task labels are used only as supervision and are not input features. DMS denotes a RibonanzaNet-derived DMS-like proxy rather than measured chemical probing. Evaluation-set sizes and split definitions are reported in Supplementary Tables S1 and S3.

| Task | Label source | Primary metric and reference | Result | Interpretation and controls |
| --- | --- | --- | --- | --- |
| Nucleotide identity | PDB sequence | Weighted recovery; random 25% | 64.6% [63.7, 65.6]% | Defining confidence output; direct sequence information in local tertiary geometry |
| Base-pair state | DSSR paired/unpaired label | Accuracy and AUC; majority 63.1% | 79.2%; AUC 0.880 | Confidence association AUC $-0.5 = -0.01$; no DSSR, dot-bracket or pair labels used as features |
| DMS-like proxy | RibonanzaNet-inferred reactivity | Pearson $r$ and $R^2$; null $r = 0$ | $r = 0.576$; $R^2 = 0.329$ | Confidence association $r = +0.03$; intermediate accessibility/chemistry-associated layer, not a measured DMS predictor |
| Protein context | RNA-protein distance $\leq 6$ Å | AUC; random 0.5 | AUC 0.673 | Confidence association $= -0.03$; weak external-environment signal, with no protein atom coordinates used as features |

FORGE nucleotide confidence was nearly orthogonal to these other labels: it did not predict base-pair state (AUC 0.49), correlated weakly with the DMS-like proxy (*r* = 0.03) and was weakly anticorrelated with protein proximity (*r* =*−*0.03). Confidence is therefore not a generic proxy for whether a position is paired, exposed or protein-bound; it measures whether the local tertiary geometry constrains base identity.

### 1.6 Compatibility, foldability and design score are separable

FORGE cross-entropy (CE) scores how compatible a candidate sequence is with a fixed backbone. For 50 post-2025 chains, native sequences ranked above composition-preserving and secondary-structure-context shuffles with near-perfect AUC (0.99– 1.00), above gRNAde and RiboDiffusion designs (AUC 0.98) and above compensatory pair-swap mutants (AUC 0.89) (Fig. 5a). RDesign was an instructive exception: among 39 evaluable chains the native-above-decoy AUC was 0.41, meaning that RDesign sequences often received lower FORGE CE than the native sequence. A designed sequence can therefore score favourably under one geometric compatibility criterion without being more native, functional or experimentally successful.

**Fig. 5.**
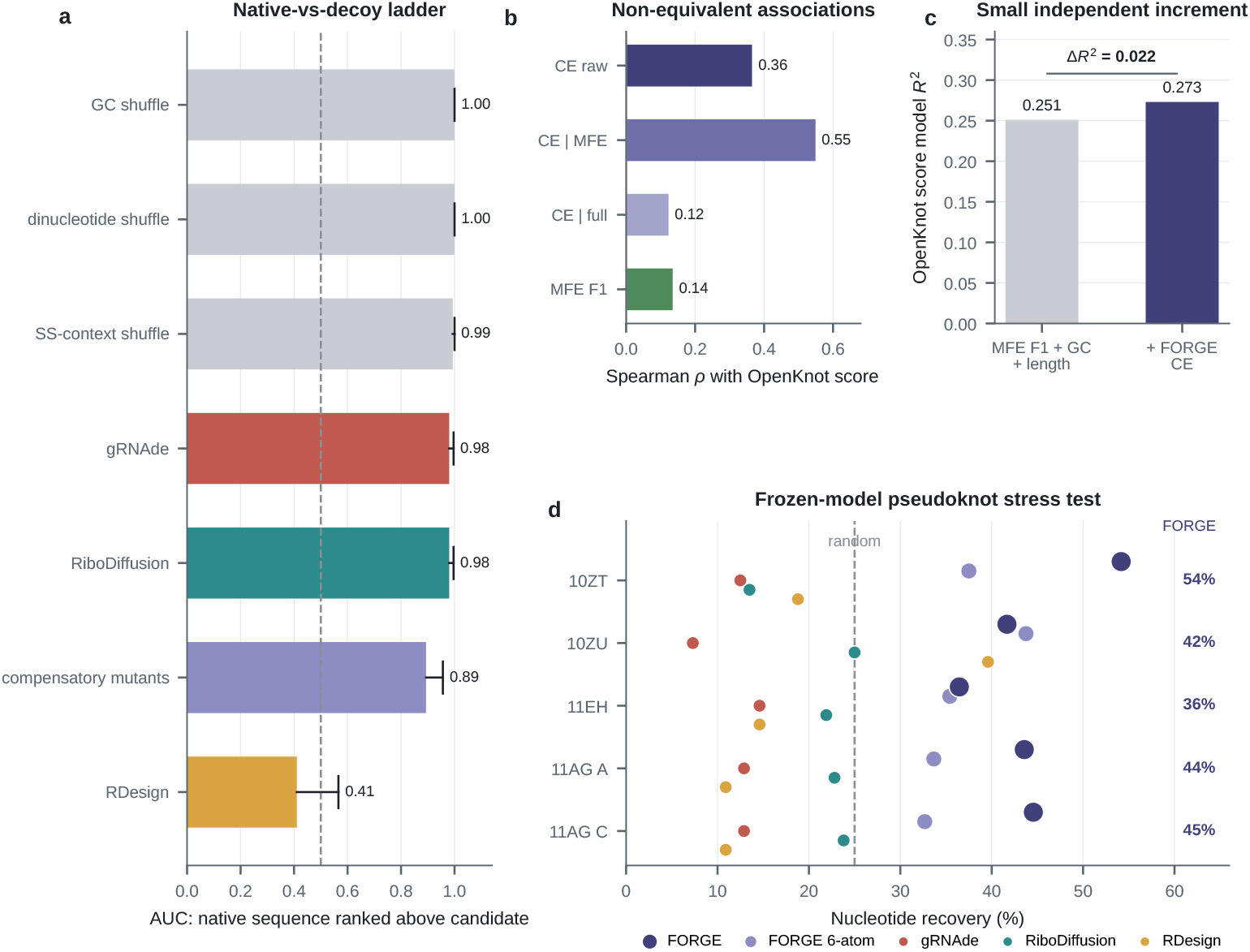
Design-objective separability and frozen-model stress tests. **a**, Native-versus-decoy AUC ladder across composition-preserving shuffles, learned inverse-folding outputs, compensatory pair swaps and RDesign outputs (error bars, Wilson 95% intervals from stored tier aggregates; dashed line, no native preference). **b**, Raw and partial Spearman associations between FORGE cross-entropy, RNAfold MFE F1 and OpenKnot experimental score in the complete-case design subset (*n* = 1, 656). **c**, Linear-model fit to OpenKnot score before and after adding FORGE cross-entropy to MFE F1, GC content and length (Δ*R*^2^ = 0.022, *p* = 4.3 *×* 10^*−*12^). **d**, Per-chain nucleotide recovery (dot plot) for FORGE, the six-atom N-free retrained model and public pretrained baselines on five solved AI-designed pseudoknot chains; the dashed line marks random recovery.

OpenKnot design records provided an independent test of the same principle. Among 8,125 pseudoknot designs, FORGE CE was positively associated with the experimental SHAPE-based score, but more weakly than RNAfold MFE F1. In the 1,656-design complete-case subset, controlling for MFE F1 raised the partial Spearman correlation to *ρ* = +0.549, and after also controlling for GC content and length it fell to *ρ* = +0.123; adding FORGE CE to a linear model of MFE F1, GC content and length contributed a small but measurable increment (Δ*R*^2^ = 0.022, *p* = 4.3 *×* 10^*−*12^; Fig. 5b,c). The positive CE coefficient is consistent with some designs departing from natural sequence–backbone preferences while still scoring well experimentally. Native-like compatibility, thermodynamic foldability and experimental design score are related but distinct objectives.

### 1.7 Solved AI-designed pseudoknots test frozen-model generalisation

To test whether the diagnostic signal generalises to non-natural backbones, we applied the frozen model to cryo-EM structures from the OpenKnot AI pseudoknot-design study [11]: PDB entries 10ZT (gRNAde), 10ZU (MPNN-fixbb), 11EH (Struct2Seq) and 11AG (MPNN-RFdiff, two chains), downloaded on 2026-05-24 and processed with the locked checkpoint. Best training-set sequence identities were 34–39%, below the prespecified threshold (Supplementary Table S2). FORGE retained above-random recovery on all five chains (54.2%, 41.7%, 36.5%, 43.6% and 44.6%; Fig. 5d), while public pretrained baselines were at or below random (gRNAde 7–15%, RiboDiffusion 13–25%, RDesign 11–40%). On 10ZT, designed by gRNAde, gRNAde itself recovered 12.5% in standard inference mode, whereas FORGE read 54.2% from the same backbone coordinates: a generative model can produce a successful fold without an invertible diagnostic readout of nucleotide identifiability.

Confidence behaviour matched the main calibration analysis. Mean per-position confidence was 0.43–0.46 on these structures (versus 0.48 on the full post-2025 set), only 1–7% of positions exceeded the 0.7 high-confidence threshold, and 13 of those 18 high-confidence calls (72%) were correct (Supplementary Fig. S4a,b). The model thus recognises out-of-distribution geometry and reduces confidence rather than hallucinating certainty. The N-free control was heterogeneous across design strategies: recovery on 10ZT dropped by 16.7 percentage points (54.2% to 37.5%) and on 11AG by 10–12 points, whereas 10ZU and 11EH were essentially unaffected, suggesting that different design methods encode the glycosidic-anchor position differently. Purine/pyrimidine accuracy remained above chance on all chains (7-atom, 65–72%; N-free, 50–62%; random 50%; Supplementary Fig. S4c).

A complementary forward-folding control quantified the reverse direction, from sequence to structure. We submitted the scaffold-embedded constructs to AlphaFold 3 [12] with three sequence classes per construct: the true design sequence, the FORGE-derived sequence and the FORGE-derived sequence after R-Net-guided reactivity refinement (Methods). Only the true 10ZT sequence returned a near-experimental fold (6.6 Å P-RMSD; all 38 native Watson–Crick pairs recovered). The remaining true sequences failed at 14–22 Å despite mostly correct base pairing (SS *F*_1_ 0.69–0.88), and the FORGE-derived and refined sequences were uniformly non-foldable (SS *F*_1_ ≤ 0.36; Supplementary Figs. S1 and S2). Read-back from geometry therefore does not invert into foldability.

## 2 Discussion

FORGE reframes RNA backbone coordinates as a measurable source of task-specific structural information rather than only an input to inverse folding. The central finding is that a seven-atom phosphate–sugar and glycosidic-anchor representation contains substantial nucleotide-identifying signal, but not enough to determine every position uniquely. This changes how sequence recovery should be interpreted: high recovery indicates a geometrically constrained nucleotide, whereas low recovery may reflect model error, missing chemical detail or genuine ambiguity in the representation.

The task-specific analyses place different annotations at different distances from the backbone. Base-pair state was readily inferred, the DMS-like proxy was captured at intermediate strength and protein-proximal context was weaker but detectable. We do not claim that FORGE predicts reactivity or binding; these readouts serve as information-distance calibration, quantifying how far each biological annotation lies from the coordinate representation. The primary signal is the ordering itself: direct conformational labels sit close to the geometry, whereas labels depending on nucleotide chemistry, solvent exposure or a binding partner sit progressively farther away. Nucleotide confidence was nearly orthogonal to all three auxiliary labels, so it should be read as a specific estimate of whether tertiary geometry constrains base identity, not as a generic measure of whether a position is structured or functionally engaged.

The representation controls locate where the nucleotide information resides. Removing the glycosidic nitrogen coordinate before retraining retained most of the discrimination, showing that phosphate–sugar geometry alone carries substantial signal, while the remaining drop shows that N1/N9 still contributes orientation information. The seven-atom representation is nevertheless a coarse model of RNA chemistry: it omits base-edge atoms, modifications, solvent, ligands, proteins and conformational ensembles. High-cross-entropy positions and mixed feature-space neighbourhoods should therefore be read as positions requiring additional evidence, not as positions where a larger classifier is guaranteed to succeed.

These properties define the practical use case. FORGE can flag high-confidence residue identities, expose ambiguous positions for manual inspection, score candidate sequences against a fixed backbone and audit inverse-folding outputs. Low-confidence positions are not failures but positions where the coarse representation is information-limited, and a conservative workflow would combine FORGE with full-atom refinement, density inspection, secondary-structure models [13], evolutionary covariation and chemical probing [8], using low-confidence predictions to focus rather than replace expert review. The per-chain abstention analysis shows that many chains contain large geometry-underdetermined regions, so this triage role is the realistic operating mode rather than a fallback.

The design analyses argue against using any single metric as a proxy for all RNA design objectives. FORGE ranks most native sequences above decoys, yet the RDesign exception and the OpenKnot analyses show that native-like compatibility, thermodynamic foldability and experimental design score are separable. The AI-designed pseudoknot stress test reaches the same conclusion from the opposite direction: a generative model can produce an experimentally resolved structure while standard inverse-folding inference does not recover the designed sequence from that backbone. The forward-folding control closes the loop between the two directions of the structure–sequence mapping. Identifiability (structure to sequence) and foldability (sequence to structure) are logically inverse problems, yet they behaved as independent objectives: AlphaFold 3 recovered the experimental fold for only one of the four solved constructs [12], and none of the sequences FORGE read out of the geometry. Two consequences follow for machine-learning practice. Sequence recovery, a standard inverse-folding metric, measures information content in the geometry and cannot be read as evidence that a recovered sequence would refold to the source structure. Conversely, folding failure does not imply absent information, because the same constructs retained 36–54% readable nucleotides in their backbone geometry. The mapping is informative in one direction and unreliable in the other, and audit metrics should be selected accordingly.

Several limitations bound these claims. The public baseline comparison uses pretrained models and is confounded by input representation, training data and architecture. The N-free control is a separate training run, so some of the difference could reflect optimisation rather than atom content. The calibration subset is easier than the full temporal set, and some family estimates and the five-chain stress test have small sample sizes. The work is entirely computational and claims no wet-lab validation or sequence-design success. Within these boundaries, FORGE provides a reproducible audit layer for RNA structural interpretation: it identifies where backbone geometry already contains annotation-relevant information and flags where base-edge chemistry, evolutionary covariation, ensemble structure or experimental evidence is required.

## 3 Methods

### 3.1 Data sources and temporal split

RNA structures and metadata were obtained from the Protein Data Bank (PDB) [1]. RNA chains were parsed from mmCIF/PDB files, and chains with sufficient coordinate coverage for the required atom set were retained. The locked manifest contains 13,251 pre-2025 training chains, 3,293 validation chains, 2,860 pre-2025 cluster-held-out test chains and 4,135 chains deposited from 1 January 2025 through the local data snapshot used for this submission. The post-2025 set was used as the primary temporal out-of-distribution evaluation. Sequence clustering was performed with CD-HIT to reduce homology leakage within the pre-2025 split [14]. Rfam family labels, when present, were used only for stratified analysis and not as model inputs [15].

### 3.2 Backbone representation and feature construction

Each nucleotide was represented by seven atoms: P, O5′, C5′, C4′, C3′, O3′ and the glycosidic nitrogen, defined as N1 for pyrimidines and N9 for purines. This representation captures phosphate-sugar geometry and an anchor towards the base, but does not include base-edge atoms such as A-N6, G-O6, C-N4 or U-O4. A *k* = 32 nearest-neighbour graph was constructed over residue centroids. FORGE computed 211 node features, including local distances, angles, dihedrals, sugar geometry, helical parameters, curvature, graph statistics and local density descriptors, and 362 edge features that were pooled by mean and standard deviation to produce 724 edge-derived features. The final vector therefore contained 935 features per nucleotide. A feature ontology was used for interpretability analyses; 889 of 935 dimensions were assigned to named categories in the current mapping.

### 3.3 Model training

The primary FORGE classifier was XGBoost with multi-class log-loss optimisation [9]. The locked training configuration used 500 estimators, maximum depth 8, learning rate 0.1, subsample 0.8 and colsample_bytree 0.8. Predictions are four-class probabilities over A, C, G and U. The confidence score is the maximum predicted probability, and sequence-level compatibility is the mean per-position cross-entropy of a candidate sequence under the predicted distribution.

### 3.4 Baselines and comparison scope

Public gRNAde, RiboDiffusion and RDesign inference pipelines were evaluated on the post-2025 structures when their input requirements were satisfied [4–6]. RDesign was evaluable on 3,966 chains in the aggregate comparison; gRNAde and RiboDiffusion were evaluable on 4,135 chains. These baselines were not retrained on the FORGE split, so the comparison does not isolate architecture, training data or input atom set. A two-layer MLP trained on the same 935 FORGE features was used as a same-representation architecture control.

### 3.5 Representation controls

For the N-removed retraining control, the glycosidic nitrogen coordinate channel was zeroed before feature extraction, and a classifier was retrained from scratch on the full pre-2025 training set (13,251 chains) with the same hyperparameters. The resulting model was evaluated on the post-2025 temporal test set (4,167 chains, 298,662 positions) and compared with the seven-atom baseline. The N-free set is larger than the main seven-atom temporal test set (4,135 chains) because the reduced atom-coverage requirement (no glycosidic nitrogen) allowed 32 additional chains to be retained. Unless otherwise stated, the main-text comparisons use the 4,135-chain seven-atom manifest. For inference-time atom-feature masking, features whose names depended on removed atoms were zeroed and the original model was applied to the full post-2025 set. The 5-atom mask removed O5′ and O3′-dependent features; the 3-atom mask retained only features compatible with P, C4′ and N. These masks were used as stress tests rather than retrained reduced-atom models.

### 3.6 Calibration and abstention

Expected calibration error (ECE) was computed with 10 confidence bins on the post-2025 calibration-evaluable subset [16]. Pooled abstention curves were generated by sorting all positions by model confidence and removing the least-confident fraction.

Per-chain abstention was computed independently for each chain as the fraction of positions that must be removed before the retained positions reach 90% accuracy. The subset used for this analysis contained 2,584 chains and 143,743 nucleotide positions. For the full temporal chain-level distribution in Fig. 4d, per-position predictions from 4,135 post-2025 chains were grouped by chain, sorted by confidence and evaluated at 10% abstention increments. Chains that did not reach 90% retained-position accuracy after removing 90% of positions were placed in the *>* 90 bin. The confidence histogram in Fig. 4c uses the same full post-2025 per-position predictions.

### 3.7 Native-vs-decoy compatibility analysis

For 50 post-2025 chains, native sequences were compared with decoys grouped into GC-preserving shuffle, dinucleotide shuffle, secondary-structure context shuffle, compensatory pair-swap and generated-design tiers. Generated-design tiers used available gRNAde, RiboDiffusion and RDesign outputs for the same backbones. FORGE cross-entropy was computed for each sequence, and AUC was defined as the fraction of native-decoy pairs in which the native sequence had lower cross-entropy than the decoy. Only tier-level native-above-decoy aggregates were stored for this analysis, so intervals use Wilson intervals from the stored aggregate counts rather than a bootstrap over raw native-decoy cross-entropy pairs.

### 3.8 Task-specific information readouts

To test whether the same RNA backbone representation contains information beyond nucleotide identity, we reused the 935 FORGE descriptors in three additional task-specific models. These analyses were not used to train the nucleotide classifier.

For base-pair prediction, DSSR-derived dot-bracket annotations were converted to paired and unpaired residue labels [7]. No DSSR pairing labels, dot-bracket annotations or base-pair identities were used as input features; all 935 geometric descriptors are computed from atomic coordinates alone (Supplementary Table S3). The model was trained on the temporal training split and evaluated on post-2025 chains with retained DSSR labels. The reported metrics include accuracy, balanced accuracy, AUC, average precision, paired recall and paired precision.

For the DMS-like proxy, per-position labels were generated with the retained RibonanzaNet inference pipeline and used as computationally derived reactivity estimates. Experimental DMS-based probing reports nucleotide accessibility through dimethyl-sulfate reactivity [8], but the labels used here are a computational proxy and should not be interpreted as experimental DMS measurements. No new wet-lab reactivity data were generated for this study. An XGBoost regressor with 300 trees and maximum depth 6 was trained on the pre-2025 training split and evaluated on the post-2025 test split. The retained evaluation arrays contain 299,074 test positions. We report *R*^2^, Pearson correlation, Spearman correlation, mean absolute error and root mean squared error.

For protein-proximal context prediction, RNA residues in protein–RNA complexes were labelled by distance to protein atoms. The primary reported setting used a 6 Å cutoff and an RBP-only post-2025 test subset; pure RNA chains were used as negative examples during training to avoid a collapsed complex-versus-free RNA classifier. Reported metrics include AUC, average precision, F1, balanced accuracy, recall for binding and non-binding residues, and binding-site precision. This task is interpreted as an indirect context layer because the RNA-only seven-atom representation does not encode protein-surface chemistry.

Confidence orthogonality was assessed by comparing the original nucleotide confidence score with base-pair state, DMS-like proxy labels and protein-proximal labels. This analysis tested whether nucleotide identifiability is merely a proxy for other structural annotations.

### 3.9 OpenKnot design analysis

OpenKnot design records were obtained from the Eterna OpenKnot benchmark data included in the local project archive. For 8,125 scored designs, RNAfold from ViennaRNA was used to compute minimum-free-energy structures and base-pair F1 scores against target structures [13]. Raw all-design associations were reported with Spearman correlation. Multivariate analyses were restricted to the 1,656-design complete-case subset specified in the locked secondary-structure evaluation summary. Partial Spearman correlations were computed by rank-transforming variables, regressing each variable of interest on the listed covariates by least squares and correlating the residual ranks. Incremental *R*^2^ was computed by ordinary least squares, comparing a base model containing MFE F1, GC content and sequence length with a full model that additionally contained FORGE cross-entropy.

### 3.10 AI-designed pseudoknot stress test

Newly deposited AI-designed pseudoknot structures from the OpenKnot study were used as a frozen-model stress test [11]. PDB entries 10ZT, 10ZU, 11EH and 11AG were downloaded from RCSB on 2026-05-24 and processed with the locked feature pipeline and model checkpoints. The analysed chains were 10ZT_A, 10ZU_A, 11EH_A, 11AG_A and 11AG_H_C. FORGE seven-atom predictions, six-atom N-free predictions and public pretrained baseline predictions were compared by per-chain recovery. Position-level confidence was analysed descriptively because the stress set contains only five chains and is too small for formal calibration. Training-set sequence redundancy was checked by global chain-level sequence alignment against the frozen training manifest; no chain exceeded the prespecified identity threshold.

### 3.11 AlphaFold 3 forward-folding control

Forward-folding predictions used the AlphaFold 3 server (alphafoldserver; ref. [12]) with default settings (five seeds per job, one seed requested; no templates or MSAs provided). Inputs were the full scaffold–design constructs as deposited (10ZT, 519 nt; 10ZU, 518 nt; 11EH, 519 nt; 11AG, homodimeric 2 *×* 518 nt), because the cryo-EM models were obtained with the design sequences embedded in the circularly permuted group II intron scaffold [11]. The designed segment (96–101 nt per chain) was extracted from the AlphaFold 3 models by sequence identity to the deposited sequence; for 11AG the homodimer was submitted as two chains and the design segments of chains A and C were evaluated independently.

Three sequence classes were submitted per construct: (i) true design sequence; (ii) FORGE-derived sequence, obtained by taking the per-position maximum-probability nucleotide of the frozen seven-atom FORGE model on the cryo-EM design segment (36.5–54.2% identity to the true design); and (iii) FORGE (R-Net optimised) sequence, obtained by gradient-free optimisation of the R-Net foundation model predictions [11] against the cryo-EM secondary structure (DMS+2A3 reactivity channels; target profile 0 for paired positions and 1 for unpaired positions; 10–16 single-nucleotide or compensating base-pair-swap mutations, high-confidence FORGE positions locked, RNAfold MFE *F*_1_ 0.56–0.98 with the target structure; the best-scoring variant is reported here).

Comparison metrics were computed on the designed segment after rigid-body fit to the cryo-EM design segment. P-RMSD is the mean backbone phosphate root-mean-square deviation; WC pair recovery (SS *F*_1_) uses geometric Watson–Crick detection (min donor–acceptor heavy-atom distance ≤4.2 Å; G:C, A:U and G:U); native tertiary contact recovery is the fraction of cryo-EM inter-nucleotide heavy-atom contacts (*<*7.5 Å, sequence separation *>* 2) reproduced by the predicted structure. PDB-code labels are used throughout for readability; the deposits contain the complete construct sequence with only the design segment modelled.

### 3.12 Feature-space and ontology analyses

The feature-space ambiguity analysis used leave-one-chain-out nearest-neighbour search in the 935-dimensional feature space. The analysis included 80 reference chains and 40 query chains from the post-2025 test set, restricted to chains of 10–200 nt. For each of 1,134 query positions, neighbours were drawn only from different chains. Reported statistics include *k* = 1, 3, 5, 10, 25 accuracy, the *k* = 10 accuracy trend across FORGE cross-entropy bins and the Spearman correlation between local label entropy and FORGE cross-entropy. Ontology-level pair discrimination was computed by aggregating pairwise F-statistics for nucleotide pairs within predefined feature categories.

### 3.13 Statistical reporting

All reported sample sizes correspond to the locked local result files. Chain-level means are used where chain independence is the relevant unit; pooled position-level analyses are labelled as such. Chain-bootstrap intervals were computed for weighted recovery when per-chain prediction artifacts were available. Several analyses still treat nucleotides within a chain as exchangeable observations and therefore may under-estimate uncertainty. Values are interpreted primarily by effect size, direction and consistency across controls. Exact confidence intervals are reported only where the necessary raw or chain-level artifacts were available; otherwise the corresponding limitation is stated in the source-data index or Supplementary Information.

## Data availability

All source RNA structures analysed in this study are publicly available from the Protein Data Bank at https://www.rcsb.org/. The local post-2025 temporal test manifest contains 4,135 RNA chains. OpenKnot design data were obtained from the Eterna OpenKnot benchmark repository (https://github.com/eternagame/OpenKnotAIDesignData). The processed data supporting the figures and tables are available in Zenodo at https://doi.org/10.5281/zenodo.22764225. The record contains `forge_paper_source_data_v0.1.0.tar.gz`, comprising figure source-data files, locked result JSON/CSV/NPY artifacts, baseline predictions, OpenKnot score tables and stress-test per-position outputs, and `forge_public_input_manifests_v0.1.0.tar.gz`, comprising PDB chain manifests, sequence files, split tables and small Rfam/OpenKnot-derived inputs used to identify the reused public datasets. The same record includes a file-level manifest with SHA-256 checksums and a manuscript source-data map linking each main and supplementary figure or table to its underlying data artifacts.

## Code availability

The public release includes the FORGE package, training scripts, inference scripts, baseline wrappers, figure-generation scripts, trained XGBoost models, environment specification, locked result manifests and the exact scripts used to generate the figure source-data files in the Zenodo archive `forge_code_and_models_v0.1.0.tar.gz`: https://doi.org/10.5281/zenodo.22764225. The command-line interface exposes `forge featurize, forge predict` and `forge explain`.

## Materials availability

No new biological materials were generated in this computational study.

## Acknowledgements

We thank the Eterna community for releasing OpenKnot design data and the developers of the PDB, Rfam, ViennaRNA, XGBoost, DSSR and related open-source tools. This work received no external funding.

## Author contributions

L.G., J.L., X.T. and B.L. conceived the study and jointly designed the model. L.G. and B.L. designed the model-evaluation experiments. L.G. implemented the computational analyses, prepared the figures and drafted the manuscript. All authors interpreted the results, revised the manuscript and approved the final version.

