## Supplementary_information for "FORGE audits residue-level information encoded in RNA tertiary-structure geometry"

#### Supplementary Note S1: AlphaFold 3 forward-folding control

The forward-folding control evaluates whether the structure-to-sequence information read by FORGE can be inverted by a contemporary folding model. The four solved AI-designed pseudoknot constructs (PDB 10ZT, 10ZU, 11EH and 11AG; design sequences embedded in a circularly permuted group II intron scaffold) were submitted to the AlphaFold 3 server as the full deposited constructs, matching the cryo-EM sample composition (11AG as a two-chain homodimer). Three sequence classes were compared per construct: the true design sequence, the FORGE-derived sequence (per-position maximum of the frozen-model readouts, 36.5–54.2% identical to the true design) and the FORGE-derived sequence after R-Net-guided reactivity refinement (48–59 mutations; reactivity profiles optimised against the cryo-EM secondary structure with FORGE high-confidence positions locked).

Evaluation used the designed segment (96–101 nt) after rigid-body fit to the cryo-EM model. P-RMSD is the mean backbone-phosphate deviation; SS $F_1$ is the $F_1$ of geometric Watson–Crick pairs (donor–acceptor distance ≤4.2 Å); contact recovery is the fraction of cryo-EM inter-nucleotide contacts (<7.5 Å) reproduced. Five seeds were requested per job; the best-scoring model per job is reported.

Only the true 10ZT sequence returned a near-experimental fold (6.6 Å P-RMSD, SS $F_1$ 0.99, 38/38 native pairs recovered). The remaining true sequences failed at 14.1–21.9 Å despite high secondary-structure recovery (SS $F_1$ 0.69–0.88;

register shifts within two base pairs recover 32–34 of 32–35 pairs), demonstrating secondary-structure recovery without tertiary-geometry recovery. The FORGE-derived sequences were the worst input class (SS $F_1 \leq 0.13$ on three of four constructs), and R-Net-guided refinement changed foldability only marginally (SS $F_1$ 0.00–0.36; the partial exception 11AG improved from 0.13 to 0.36 at 17.3 Å P-RMSD). Sequence recovery read out of geometry therefore does not invert to foldability, with or without reactivity-guided sequence optimisation.

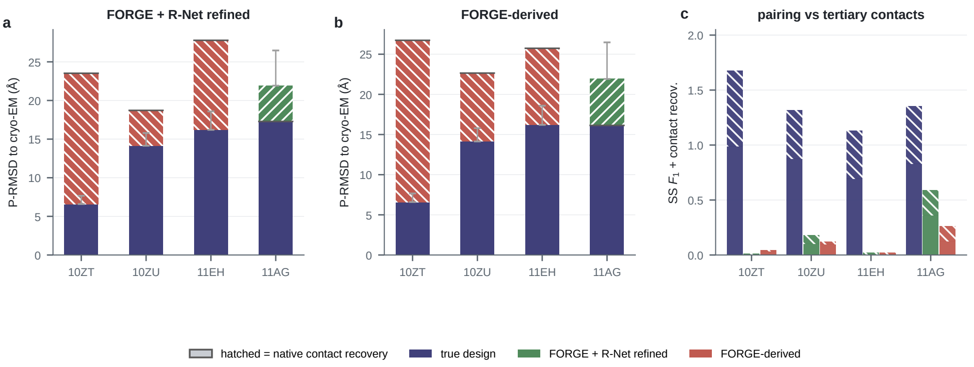


**Fig. S1 Forward-folding evaluation of the AI-designed pseudoknot constructs (AlphaFold 3). a,** Delta P-RMSD of the FORGE-derived sequence after R-Net-guided refinement relative to the true design sequence (hatched segments: green, improvement; red, degradation; grey error bars, five-seed range of the true sequence). **b,** Same for the FORGE-derived sequence. **c,** Watson–Crick pair recovery (SS $F_1$, solid) and native tertiary contact recovery (hatched) for the three sequence classes.

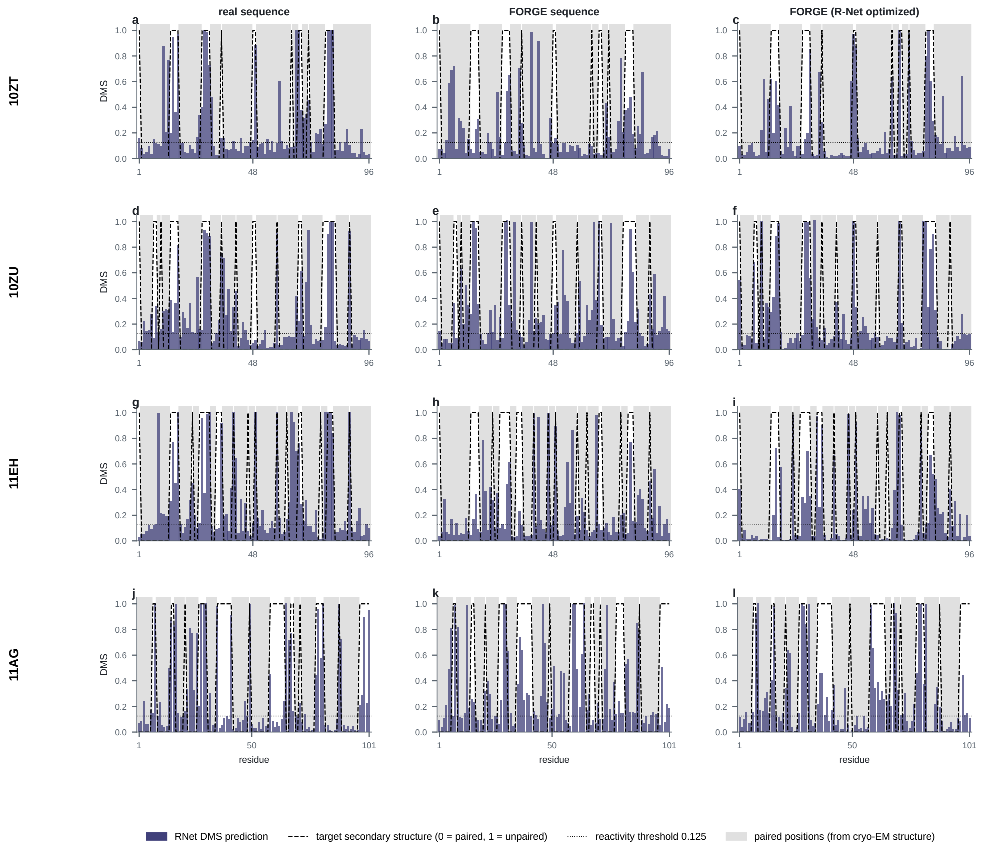


**Fig. S2 R-Net reactivity predictions against the target secondary structure.** Rows, PDB constructs; columns, sequence classes (true design, FORGE-derived, FORGE-derived after R-Net refinement). Grey bands mark positions paired in the cryo-EM structure; the dashed line is the binary target profile (0, paired; 1, unpaired); the dotted line is the reactivity threshold 0.125.

### Supplementary Table S1: Evaluation subsets

| Analysis | Chains | Positions | Selection Criterion | Metric Type | Key Value |
|---|---|---|---|---|---|
| Full post-2025 OOD | 4,135 | 291,331 | All post-2025 RNA chains | Weighted (micro) | 64.6% [63.7, 65.6] |
| Full post-2025 OOD (per-chain) | 4,135 | 291,331 | All post-2025 RNA chains | Per-chain mean (macro) | 63.4% [62.5, 63.9] |
| Calibration subset | 2,584 | 143,743 | Feature extraction succeeded | Pooled (micro-avg) | 73.1% |
| Calibration per-chain | 2,584 | 143,743 | Same as above | Per-chain mean (macro) | 63.4% [62.7, 64.1] |
| N-removed retrain (full) | 4,167 | 298,662 | All post-2025, N coord zeroed | Weighted (micro) | 58.5% [57.4, 59.6] |
| N-removed retrain (matched) | — | 72,787 | Subsampled 50% of test | Position (micro) | 66.7% [66.3, 67.0] |
| Native-vs-decoy | 50 | — | 20–200 nt, baseline designs available | Chain-level AUROC | 0.41–1.00 by tier |
| Stress test | 5 chains | 490 | AI-designed pseudoknots, cryo-EM | Per-chain | 36.5–54.2% |
| $k$-NN feature-space | 40 query | 1,134 | 10–200 nt, ref pool 80 chains | Position (micro) | $k$=10: 53.6% |
| OpenKnot complete-case | 1,656 | — | MFE F1 available | Design-level $\rho$ | $\rho_{\text{partial}}$=+0.549 |

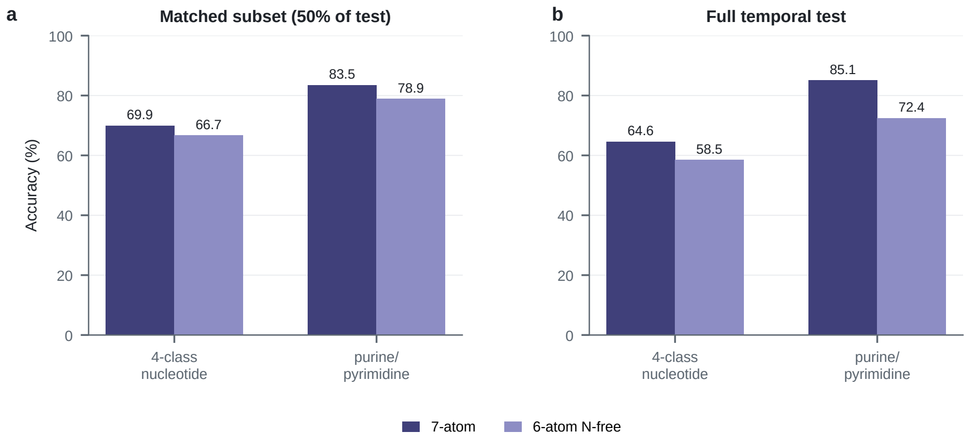


**Fig. S3 N-free retraining controls at both evaluation scales. a,** Matched-subset retraining (200,000 training positions, 72,787 test positions): the seven-atom model reached 69.9% four-class recovery and 83.5% purine/pyrimidine accuracy, and the six-atom N-free model 66.7% and 78.9%. **b,** Full temporal retraining on 13,251 pre-2025 chains and the full post-2025 test set (4,167 chains, 298,662 positions): the seven-atom model reached 64.6% weighted recovery and 85.1% purine/pyrimidine accuracy, and the N-free model 58.5% and 72.4%. Both scales show that most nucleotide-discriminating signal persists without the glycosidic nitrogen coordinate.

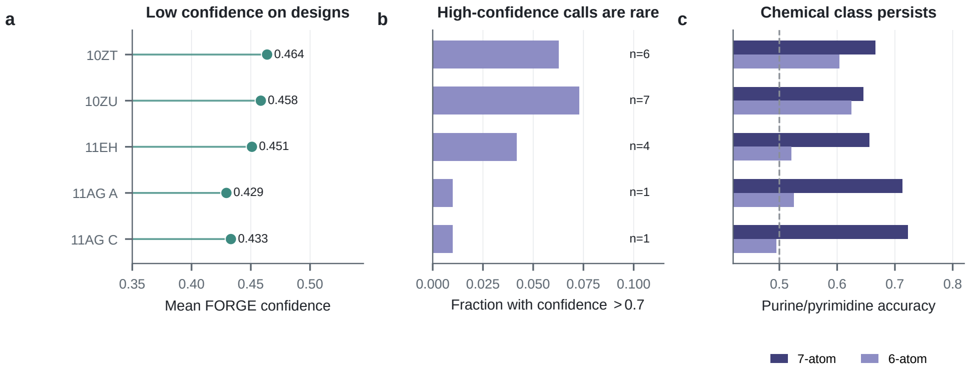


**Fig. S4 Confidence and chemical-class behaviour on the solved AI-designed pseudoknots. a,** Mean per-position FORGE confidence per chain (full post-2025 reference, 0.48). **b,** Fraction of positions with confidence >0.7, annotated with the absolute number of high-confidence positions. **c,** Purine/pyrimidine accuracy for the seven-atom and six-atom N-free models (dashed line, random 50%).

### Supplementary Table S2: Stress-test chain homology verification

| Chain | Length | In training? | Best seq. identity | Best match | PDB deposit | Download date |
|---|---|---|---|---|---|---|
| 10ZT (gRNAde) | 96 | No | 36.0% | 5O9Z_X_2 (100 nt) | 2026-05-13 | 2026-05-24 |
| 10ZU (MPNN-fixbb) | 96 | No | 38.5% | 3ADC_H_C (88 nt) | 2026-05-13 | 2026-05-24 |
| 11EH (Struct2Seq) | 96 | No | 34.4% | 3A2K_H_C (77 nt) | 2026-05-14 | 2026-05-24 |
| 11AG A (MPNN-RFdiff) | 101 | No | 35.6% | 6CK4_H_D (101 nt) | 2026-05-14 | 2026-05-24 |
| 11AG C (MPNN-RFdiff) | 101 | No | 35.6% | 6CK4_H_D (101 nt) | 2026-05-14 | 2026-05-24 |

### Supplementary Table S3: Multi-task label provenance and leakage control

| Task | Label source | Forbidden inputs removed | Split | Baseline |
|---|---|---|---|---|
| Nucleotide identity | PDB sequence | None; 7-atom coordinates only | Pre-2025 / post-2025 temporal | 25% random |
| Base-pair state | DSSR dot-bracket | No DSSR, dot-bracket or pair labels used as features; all 935 descriptors computed from coordinates alone | Pre-2025 / post-2025 temporal | 63.1% majority class |
| DMS-like proxy | RibonanzaNet inference | Sequence not used as input; backbone coordinates only | Pre-2025 / post-2025 temporal | Mean predictor |
| Protein-proximal context | PDB inter-atomic distance $\leq 6$ Å | No protein atom coordinates used as input | Pre-2025 train / post-2025 RBP-only test | Class-frequency baseline |
